# Validation of bronchial airway gene expression associated with bronchiectasis in nasal epithelium

**DOI:** 10.1101/2025.02.26.639889

**Authors:** Jason Wong, Whitney Souery, Alejandro A. Diaz, Ke Xu, Xiaohui Xiao, Hanqiao Liu, Gang Liu, Adam C. Gower, Yuriy O. Alekseyev, Ehab Billatos, Marc E. Lenburg, the DECAMP investigators

## Abstract

**Objectives:** Examination of bronchial epithelium-derived gene expression signature of bronchiectasis (BE) in nasal epithelium.

**Methods:** We studied 220 participants from the Detection of Early Lung Cancer Among Military Personnel study with bulk RNA-seq of nasal epithelium brushings. Gene set enrichment analysis (GSEA) was used to examine whether genes previously identified as increased or decreased in the bronchial epithelium of individuals with radiologic BE are significantly enriched among the genes most significantly altered in the nasal epithelium. GSEA and cell-type specific gene expression signatures were used to examine changes in the cellular composition of nasal epithelium associated with radiologic BE.

**Results:** Genes previously found to have significantly altered expression in bronchial epithelium in radiologic BE are concordantly enriched among the genes most increased or decreased in nasal epithelium in radiologic BE in this validation set (p = 3×10^−7^, increased; p = 6×10^−8^, decreased). GSEA performed using cell-type signatures demonstrates that in nasal epithelium, BE is associated with increased multiciliated and deuterosomal cell-related gene expression and decreased expression of basal cell-related gene expression, consistent with our previous findings in bronchial epithelium.

**Conclusion:** Our work validates our bronchial signature of BE in an independent dataset and demonstrates that similar gene expression changes are associated with radiologic BE in the bronchial and nasal epithelium. These findings support considering nasal epithelium brushing as a less invasive tool for screening and monitoring BE.

**Summary:** This study validates a previously reported bronchial signature of bronchiectasis (BE) in nasal epithelium. Changes in gene expression and cell-type composition associated with radiologic BE are similar across both bronchial and nasal epithelium.

## Introduction

Bronchiectasis is considered the final common pathway for numerous diseases, including rheumatologic conditions, primary ciliary dyskinesia, cystic fibrosis, and pulmonary infections.^1^ It is also associated with chronic obstructive pulmonary disease (COPD).^2-5^ Patients with BE experience chronic cough, sputum production, dyspnea, and hemoptysis.^6^ They also have frequent exacerbations and chronic infection, significantly increasing their risk of morbidity and mortality.^7^ On computed tomography (CT), BE is associated with imaging features, including an airway-to-arterial ratio > 1, a lack of tapering of bronchi, and airway visibility within 1 cm of a costal pleural surface or touching the mediastinal pleura.^8-10^ Treatment of BE involves addressing the underlying etiology, reducing infection risk with antibiotics, and managing exacerbations.^11^

Transcriptomic analysis of the airway epithelium has previously been employed in cancer ^12,13^, COPD ^14,15^, emphysema ^16^, and interstitial lung abnormalities.^17^ The concept of an airway field of injury enables profiling the bronchial epithelium to assess molecular changes occurring distal to the disease site.^18^ By identifying gene expression changes associated with CT features compatible with BE (radiologic BE) in normal-appearing bronchial epithelial cells from participants without physician-diagnosed disease, we identified a 655 gene signature consisting of five co-expression clusters based on hierarchical clustering.^19^ Pathway enrichment analysis showed that clusters with decreased expression in BE were linked to cell adhesion and Wnt signaling, while those with increased expression were associated with endopeptidase activity and ciliogenesis. The gene expression pattern of the bronchial signature genes separated participants into three clusters: normal, intermediate, and bronchiectatic. The bronchiectatic cluster was enriched for participants with more lobes of radiologic BE and more BE-associated symptoms reported, as compared to the normal and intermediate participant clusters. Cell-type deconvolution suggested that radiologic BE in bronchial epithelium is associated with increased proportions of ciliated and deuterosomal cells and a decreased proportion of basal cells.

Our analysis suggests overall similarities between widespread radiologic BE and clinical BE, as supported by the prevalence of participants in the bronchiectatic subgroup meeting both radiologic and clinical criteria for BE. Moreover, in addition to describing gene expression related to biological processes previously associated with BE development (e.g., decrease in cell adhesion and increase of inflammatory processes), our results also highlighted novel mechanisms that may be associated with the initiation of BE, such as increased expression of genes involved in ciliogenesis and decreased expression of genes involved in Wnt signaling pathways.^19^

In the present study, we tested the hypothesis that BE-associated molecular changes in bronchial epithelium might extend to nasal epithelium, and sought to validate the bronchial-derived signature of BE in independent samples from the DECAMP-1 and DECAMP-2 cohorts.

## Methods

### Study participants

Study participants were drawn from the DECAMP-1 and DECAMP-2 studies. The details of the Detection of Early Lung Cancer Among Military Personnel (DECAMP) protocols were published previously.^20^ The DECAMP studies recruited from 15 military treatment facilities, Veterans Affairs hospitals and academic centers, researching molecular markers of lung cancer and lung cancer risk and collecting biospecimens such as nasal and bronchial brushings from normal-appearing airway epithelium. DECAMP-1 (NCT01785342) recruited current and former tobacco users with indeterminate pulmonary nodules (7-30 mm), while DECAMP-2 (NCT02504697) included current and former tobacco users at high risk of developing lung cancer (i.e., 20+ pack-year smoking history and either COPD or a family history of lung cancer) eligible for lung cancer screening by chest CT. None of the DECAMP participants reported physician-diagnosed bronchiectasis at the time of enrollment. Per DECAMP protocol, each participant was asked to complete a Lung Health Questionnaire covering demographics, personal medical history, family medical history, medications, smoking history, alcohol and recreational drug history, and symptom history such as cough, dyspnea and sputum production. This study was approved by the Human Research Protection Office (HRPO) for the United States Department of Defense and the institutional review board of every participating site. All participants provided written informed consent to participate in the study.

### Computed Tomography (CT) protocols

DECAMP-2 used a standardized protocol for image acquisition and reconstruction, while DECAMP-1 CT scans were acquired as part of routine clinical care. For DECAMP-2, volumetric scans were acquired with a low radiation dose helical technique on a minimum 16-slice multidetector scanner. Scans were acquired at 2.5 to 5□mm and reconstructed into 1.25□mm slice thickness using standard and high spatial frequency convolution kernels.

### Radiologic bronchiectasis ascertainment

BE was detected visually by a pulmonologist (A.A. Diaz) with more than 10 years of experience in lung imaging, blinded to the gene expression profiles and participants’ clinical data. Radiologic BE was defined with one or more of the following criteria: 1) airway dilation (airway lumen diameter greater than adjacent pulmonary vessel diameter); 2) abnormal airway tapering of any extent (no decrease in or increase in lumen moving from proximal to distal airways); and 3) visualization of a bronchus within 1 cm of the pleura. The lingula was considered a separate lobe. Widespread radiologic BE was defined as radiologic BE in ≥ 3 lobes.^21^

### RNA isolation, sequencing and data pre-processing

Nasal brushings were obtained from participants enrolled in DECAMP-1 and DECAMP- 2. Total RNA was isolated using the miRNeasy Mini Kit (Qiagen, Valencia, CA). RNA integrity was quantified by Agilent Bioanalyzer, and RNA purity was confirmed using a NanoDrop spectrophotometer. Libraries were generated using the Illumina TruSeq Stranded Total RNA kit and sequenced on Illumina NextSeq 500 instruments (with 50- or 75-base paired-end reads), Illumina HiSeq 2500 instruments (with 75-base paired-end reads), or Illumina NextSeq 2000 instruments (with 100-base paired-end reads) (Illumina, San Diego, CA) to an average read depth of 70 million reads per sample. We developed an automatic pipeline (https://github.com/compbiomed/RNA_Seq) based on the Nextflow framework to obtain the expression levels for each gene.^22^ Reads were aligned to the Genome Reference Consortium human build 37 (GRCh37) using STAR.^23^ Both gene and transcript level counts were calculated using RSEM with the Ensembl v75 annotation.^24^

### Bulk RNA-seq and sample analysis

We previously deposited nasal gene expression data from n=288 DECAMP participants in the NCBI Gene Expression Omnibus (GEO) (Series GSE210660)^25^. Of these, n=136 were from individuals who had been scored for radiologic BE who were not included in the BE signature discovery set. We identified n=132 additional participants not included in the discovery set from whom we had generated nasal RNA-seq data. We have deposited these data in GEO (submission currently pending). Of these participants, n=84 had also been scored for radiologic BE. Collectively, we refer to this combined dataset from participants not included in the BE signature discovery set who also had nasal gene expression data and radiologic BE assessment (n=220) as the nasal validation set (**Supplemental Figure 1**). The matrix of aligned counts per gene was filtered to remove lowly expressed genes that did not have a CPM > 1 in at least 10% of samples and non-protein coding genes. Surrogate variable analysis was used to correct for heterogeneity in the data.^26^ There were 17 surrogate variables, of which the first 5 were used as covariates.

Linear modeling was performed with limma^28^ (version 3.10) in R (version 3.6.0) to assess associations between each gene’s nasal expression level and the presence of widespread radiologic BE while controlling for the expression of the first 5 surrogate variables. We ranked genes by moderated t-statistic (from most increased in the nasal epithelium of individuals with widespread radiologic BE to most decreased) and examined the enrichment of different gene sets within the tails of this ranked list using Gene Set Enrichment Analysis (GSEA). Heatmaps were used to visualize gene expression data. Hierarchical clustering using the “Ward.D2” algorithm was used to identify sample clusters.

To assess changes in the cellular composition of nasal epithelium associated with radiologic BE, we used GSEA to evaluate the enrichment of sets of cell-type marker genes reported by Deprez et al.^29^ within the nasal BE ranked list described above.

## Results

### Participant demographics, pulmonary function, and imaging measurements

Of the 220 participants screened in the nasal validation set, 52 had radiologic BE in at least one lobe and 16 had widespread radiologic BE (BE in ≥ 3 lobes; **Supplementary Table 1**). Compared to the participants in the discovery set used in Xu et al.^19^ (n = 173), in the validation set, the average number of lobes with radiographic BE was lower (p < 0.001) and fewer participants had any radiologic BE reported (p < 0.001). Among patients with any radiologic BE, the average number of affected lobes was similar in the discovery and validation set (p = 0.2) (**Table 1**). Despite this, the prevalence of widespread BE in the validation set was not significantly different from the discovery set. The validation set had significantly lower rates of current smoking than the discovery set (p = 0.001). Pulmonary function was similar between the discovery and validation sets.

**Table 1.** Comparison of clinical features between participants in the nasal validation set (n = 220) and bronchial discovery set (n = 173)

|  | **Bronchial Discovery set**<br>**(N = 173)** | **Nasal Validation set**<br>**(N = 220)** | **P-value** |
| --- | --- | --- | --- |
| Any Bronchiectasis (%) | 96 (55%) | 52 (24%) | <0.001 |
| Widespread BE (>3 lobes) (%) | 20 (12%) | 16 (7.3%) | 0.14 |
| Average number of lobes with BE (mean (SD)) | 1.06 (1.16) | 0.54 (1.18) | <0.001 |
| Male (%) | 141 (82%) | 154 (70%) | 0.01 |
| Race |  |  | 0.08 |
| Asian (%) | 5 (3%) | 5 (2%) |  |
| Black (%) | 21 (12%) | 41 (19%) |  |
| Others/Unknown (%) | 18 (10%) | 11 (5%) |  |
| White (%) | 129 (75%) | 163 (74%) |  |
| Smoking Status |  |  | 0.089 |
| Current tobacco user (%) | 75 (43%) | 71 (37%) |  |
| Former tobacco user (%) | 90 (52%) | 123 (63%) |  |
| FEV1 % predicted (mean (SD)) | 71 (18) | 66 (18) | 0.090 |
| FEV1/FVC (mean (SD)) | 0.6 (0.13) | 0.59 (0.14) | 0.71 |
| Cough Yes (%) | 77 (44%) | 101 (46%) | 0.74 |
| Phlegm Yes (%) | 76 (44%) | 99 (45%) | 0.55 |
| Shortness of Breath Yes (%) | 98 (62%) | 132 (60%) | 0.29 |
| DECAMP Cohort |  |  | <0.001 |
| DECAMP1 (%) | 129 (75%) | 84 (38%) |  |
| DECAMP2 (%) | 44 (25%) | 136 (62%) |  |

### Genes significantly altered in the bronchial epithelium are among the genes most altered in the nasal epithelium

To explore the relationship between nasal and bronchial epithelial gene expression alterations in BE, we performed GSEA using gene sets from the bronchial gene signature of BE previously described by Xu et al.^19^ We observed that genes in Xu’s bronchial signature that had increased expression in individuals with widespread radiologic BE were significantly enriched among the genes most increased in the nasal epithelium of individuals with widespread radiologic BE (**Figure 1A**; p < 1×10^−6^). Similarly, genes whose expression was decreased in widespread radiologic BE in Xu’s bronchial signature were significantly enriched among genes most decreased in the nasal epithelium of individuals with widespread radiologic BE (**Figure 1B**; p < 1 x10^−7^). This pattern extended to each of the 5 gene clusters from Xu’s bronchial BE signature (**Figure 1**).

**Figure 1.**
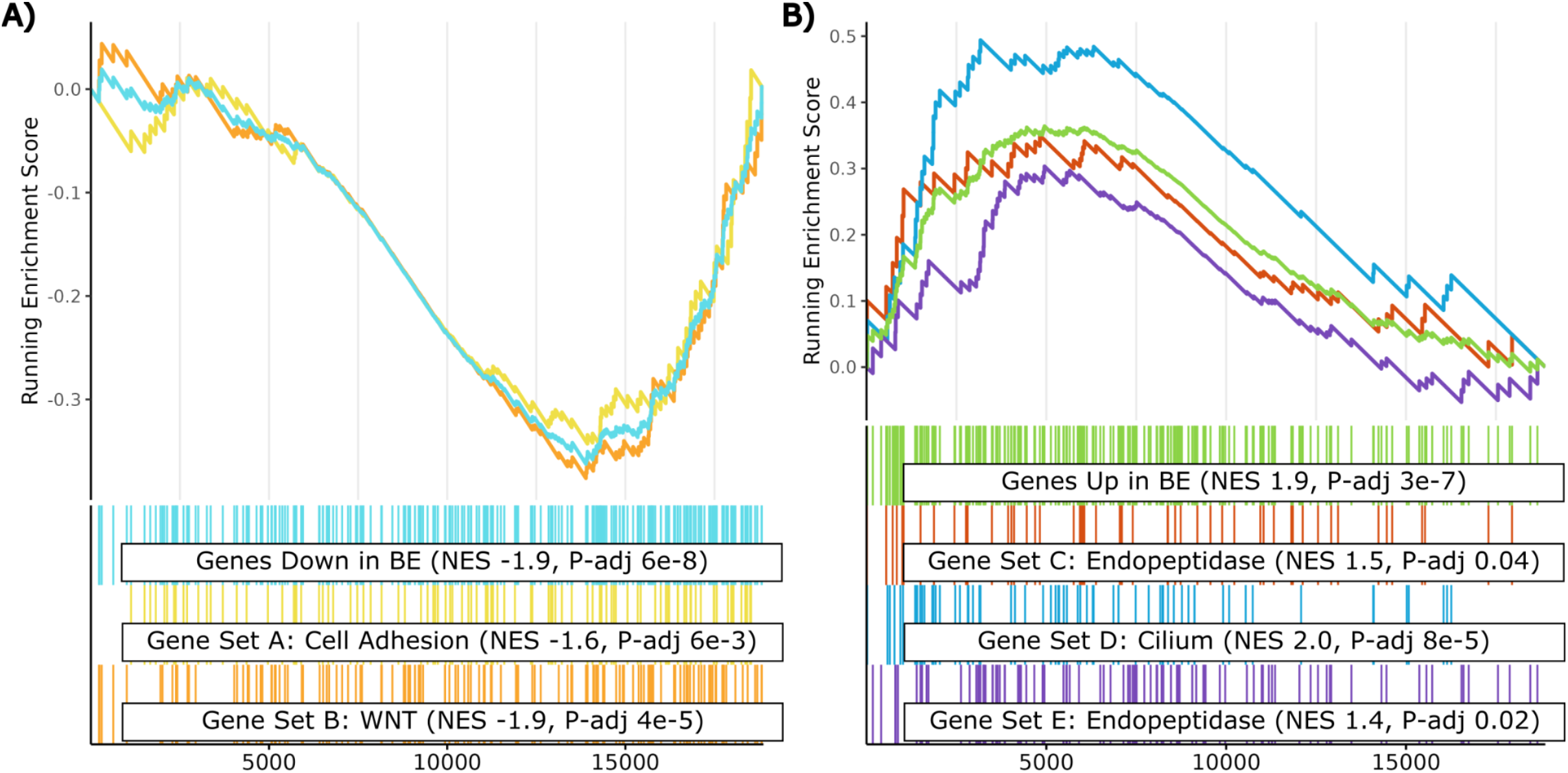
Genes with BE-associated gene expression in the bronchial epithelium are among the genes whose expression is most associated with BE in the nasal epithelium. A) Genes previously reported by Xu et al. to have decreased expression in the bronchial epithelium of individuals with widespread radiologic BE are significantly enriched among the genes most decreased in the nasal epithelium of individuals with widespread radiologic BE. This is true both for the decreased genes as a whole, as well as the two decreased gene sub-clusters identified by Xu et al. (Gene Set A and B). B) Genes previously reported by Xu et al. to have increased expression in the bronchial epithelium of individuals with widespread radiologic BE are significantly enriched among the genes most increased in the nasal epithelium of individuals with widespread radiologic BE. This is true both for the increased genes as a whole, as well as the three increased gene sub-clusters identified by Xu et al. (Gene Set C, D and E). In both plots, genes are ranked from left to right from the most positive moderated t-statistic for BE to the most negative. The vertical lines represent the position of the genes in each gene set within these ranked lists, and the y-axis represents the running GSEA enrichment score.

### Identification of two participant clusters based on gene expression profiles in nasal validation set

From the bronchial BE signature, we used the expression of the 25 genes that were the most significantly differentially expressed in either direction (up or down) with respect to widespread radiologic BE in the nasal epithelial samples to divide the research participants into two clusters(**Figure 2**). The predominant cluster (n=152) was primarily composed of patients without widespread radiologic BE. The second cluster (n=68) had a pattern of relative gene expression similar to the “bronchiectatic” cluster (increased expression of “up” genes and decreased expression of “down” genes) and included most of the patients with widespread radiologic BE. This second cluster had a higher average number of lobes with BE (p < 0.001), a higher prevalence of any radiologic BE (p < 0.001), and a higher prevalence of widespread BE (p < 0.001) (**Table 2**). The likelihood of having BE-associated symptoms such as cough, dyspnea, or phlegm did not differ significantly between the groups. Pulmonary function and smoking status were also similar between the participant clusters defined by nasal gene expression.

**Table 2.** Clinical features of participants in the nasal validation set across expression-derived clusters.

|  | **Bronchiectatic Cluster**<br>**(N = 69)** | **Normal Cluster (N = 151)** | **P-value** |
| --- | --- | --- | --- |
| Any Bronchiectasis (%) | 26 (38%) | 26 (17%) | <0.001 |
| Widespread BE (>3 lobes) (%) | 12 (17%) | 4 (2.6%) | <0.001 |
| Average number of lobes with BE (mean (SD)) | 0.99 (1.63) | 0.34 (0.83) | <0.001 |
| Male (%) | 48 (70%) | 106 (70%) | 0.64 |
| Race |  |  | 0.48 |
| Asian (%) | 1 (1%) | 4 (3%) |  |
| Black (%) | 17 (25%) | 24 (16%) |  |
| Others/Unknown (%) | 3 (4%) | 8 (5%) |  |
| White (%) | 48 (70%) | 115 (76%) |  |
| Smoking Status |  |  | 0.99 |
| Current tobacco user (%) | 23 (37%) | 48 (37%) |  |
| Former tobacco user (%) | 40 (63%) | 83 (63%) |  |
| FEV1 % predicted (mean (SD)) | 68 (18) | 65 (19) | 0.5 |
| FEV1/FVC (mean (SD)) | 0.60 (0.14) | 0.59 (0.13) | 0.7 |
| DECAMP Cohort |  |  | 0.43 |
| DECAMP1 (%) | 29 (42%) | 55 (36%) |  |
| DECAMP2 (%) | 40 (58%) | 96 (64%) |  |
| Cough Yes (%) | 30 (43%) | 71 (47%) | 0.80 |
| Phlegm Yes (%) | 27 (39%) | 72 (48%) | 0.39 |
| Shortness of Breath Yes (%) | 41 (59%) | 91 (60%) | 0.48 |
| DECAMP Cohort |  |  | 0.43 |
| DECAMP1 (%) | 29 (42%) | 55 (36%) |  |
| DECAMP2 (%) | 40 (58%) | 96 (64%) |  |

**Figure 2.**
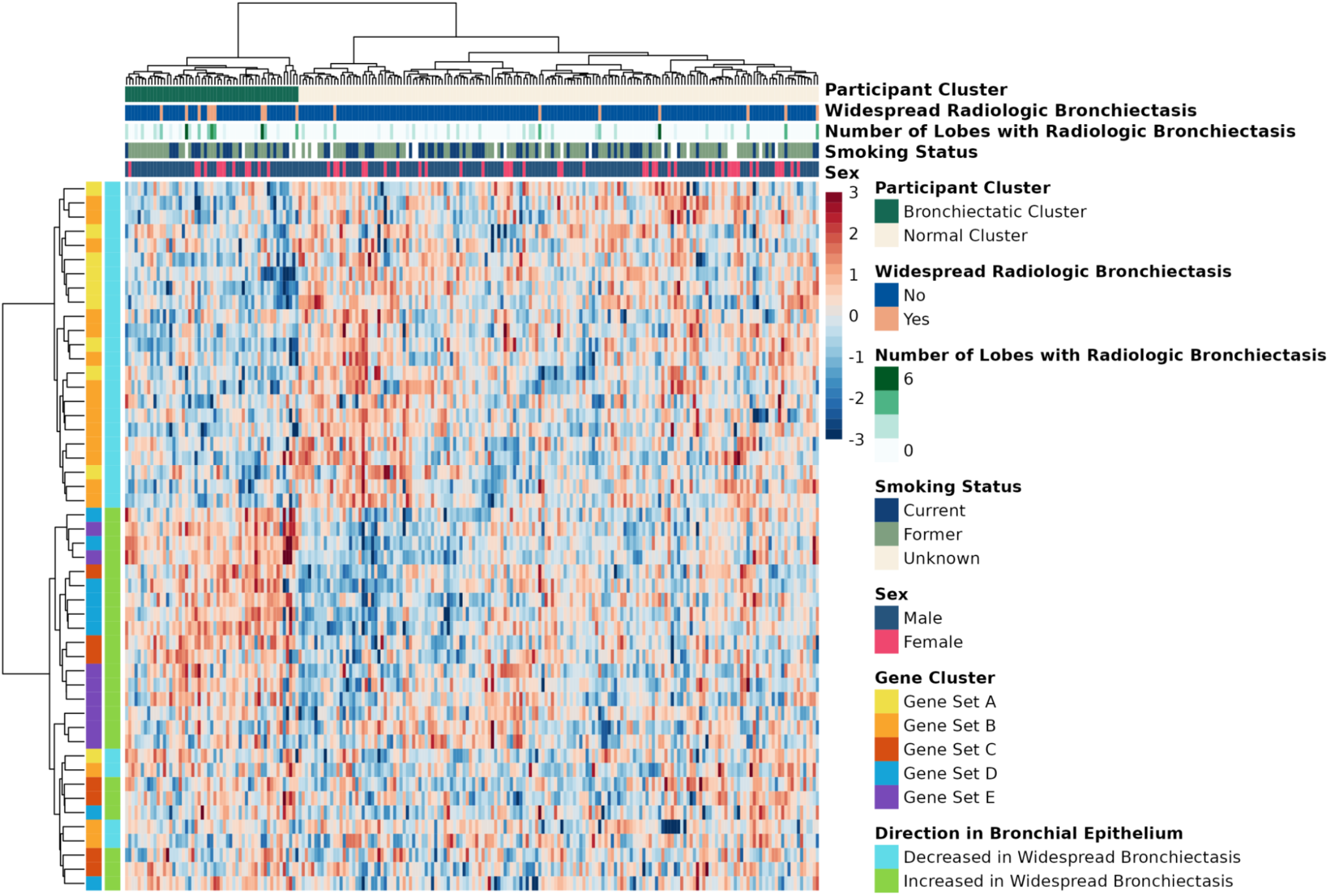
Clustering based on nasal BE-associated gene expression identifies two clusters of participants that differ with regard to radiologic BE prevalence. From the bronchial BE signature, we selected the 25 genes most significantly increased and the 25 genes most significantly decreased with respect to widespread radiologic BE in nasal epithelium to divide the research participants into two groups using hierarchical clustering. The smaller of these clusters (n = 68), which is characterized by increased expression of genes of bronchial BE “up” genes and decreased expression of bronchial BE “down” genes, is significantly enriched for participants with bronchiectasis (p < 0.001; Fisher’s Exact Test). The expression values are z-score-normalized to a mean of zero and standard deviation of one within each row, with blue, white, and red indicating z-scores of ≤ -3, 0, and ≥ 3, respectively.

### Characterizing differences in the cellular composition of nasal epithelium in bronchiectasis and as compared to bronchial epithelium

Xu et al. previously reported significant differences in the estimated cell-type proportions of basal, deuterosomal, and multiciliated cells when comparing participant clusters defined by bronchial BE signature expression.^19^ To test whether these same cell-type composition changes are present in nasal epithelium, we leveraged the same signatures from Deprez^29^ et al. for basal, deuterosomal, and multiciliated cells that were used in Xu^19^ et al. Using the nasal gene expression data, we performed GSEA to assess whether these cell-type signatures were enriched among genes most altered in the nasal epithelium of individuals with widespread radiologic BE. Deuterosomal and multiciliated cell signature genes are significantly enriched among the genes most increased in the nasal epithelium of individuals with widespread radiologic BE, while the basal cell signature genes are significantly enriched among genes most decreased (**Figure 3**).

**Figure 3.**
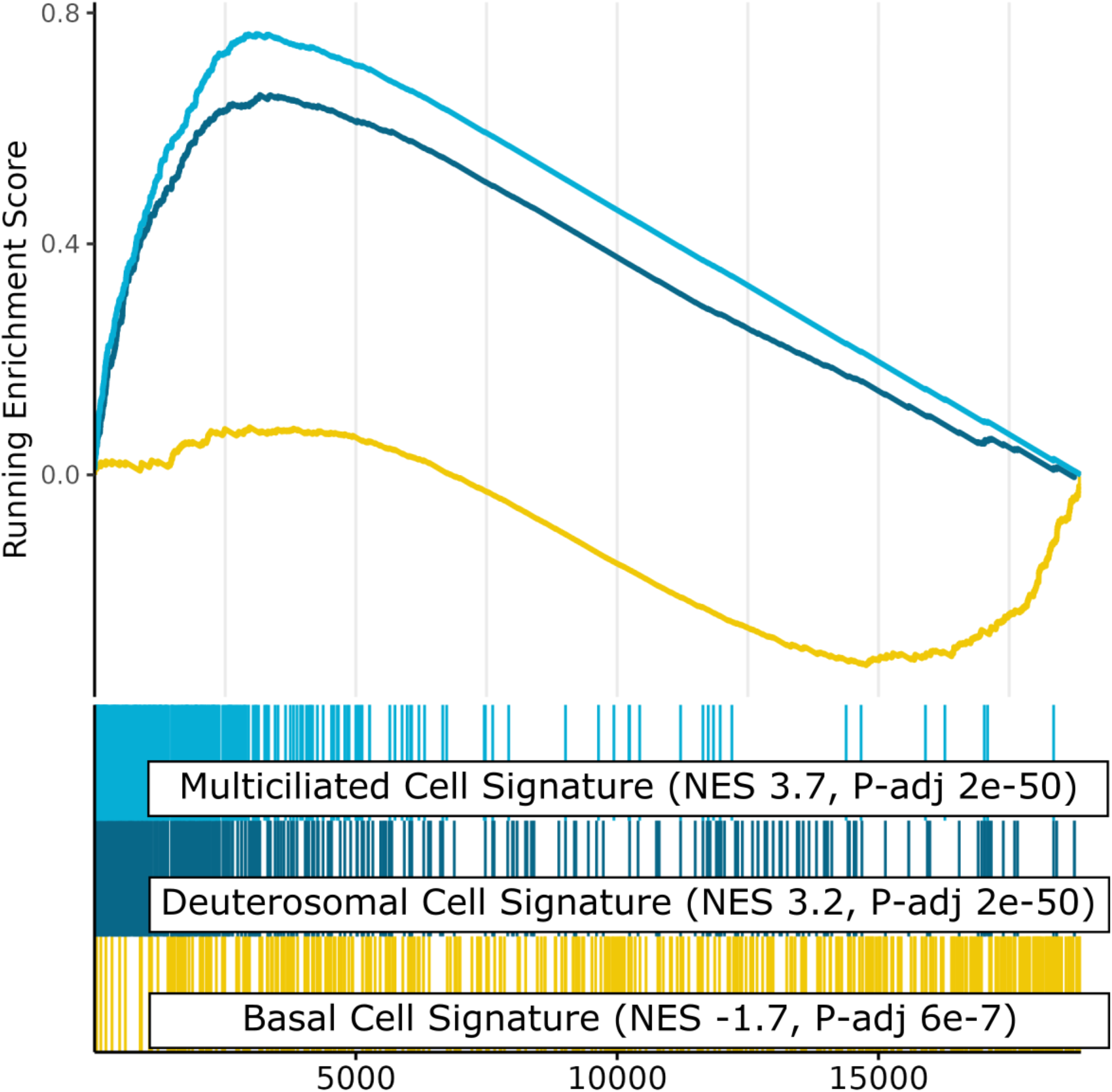
Cell-type marker gene expression suggests similar BE-associated changes in composition of bronchial and nasal epithelium. Marker genes for multi- ciliated and deuterosomal cells (as defined by Deprez^29^ et al.) are enriched among the genes whose expression is most increased in the nasal epithelium of individuals with widespread radiologic BE. In contrast, marker genes for basal cells are enriched among the genes most decreased. This is similar to the enrichment patterns seen in BE-associated bronchial epithelial gene expression and suggests that the proportion of multi-ciliated and deuterosomal cells is increased in the nasal epithelium of individuals with widespread radiologic BE while the proportion of basal cells is decreased. Genes are ranked from left to right from the most positive moderated t-statistic for BE to the most negative as in Figure 1. The vertical lines represent the position of the genes in each gene set within these ranked lists, and the y-axis represents the running GSEA enrichment score.

## Discussion

We were interested in determining the effect of BE on nasal epithelial gene expression, as these samples are non-invasive and easy to obtain. Our group previously described a signature of 655 genes whose expression is altered in the bronchial epithelium of individuals with widespread radiologic BE.^19^ We hypothesized that these transcriptomic alterations might be shared between the bronchial and nasal epithelium due to an “airway field of injury” effect.^18^

In our present study, we validated the bronchial BE signature in nasal epithelium using an independent dataset of 220 participants drawn from the same clinical cohorts as the discovery set. Compared to the bronchial discovery set, participants in the nasal validation set had similar rates of widespread radiologic BE (≥ 3 affected lobes) but significantly lower rates of any radiologic BE (≥ 1 affected lobe). We also found no association between widespread radiologic BE and shortness of breath in the validation set, in contrast to the discovery set, where these variables were positively correlated.

Our results demonstrate that the genes in the previously identified bronchial BE signature are significantly and concordantly enriched among the genes whose expression in the nasal epithelium is most strongly associated with widespread radiologic BE. While Xu et al. found 655 genes whose expression was significantly associated with widespread radiologic BE (FDR *q* < 0.1, |log_2_ fold-change| > 0.25) in bronchial epithelium in a discovery set from 173 participants, we did not detect genes whose nasal expression pattern is significantly associated with widespread radiologic BE in our larger dataset derived from 220 participants using the same criteria (data not shown). One possible explanation for our inability to identify significantly differentially expressed genes in nasal epithelium despite our significant enrichment results is that there are additional sources of gene expression variability in nasal epithelium that may make it more difficult to discover BE-related gene expression in this tissue. Alternatively, the BE-associated field of injury might be stronger in the bronchial epithelium. Smoking-associated gene expression changes have also been observed to be more robust in bronchial than in nasal epithelium.^30^

The participants were divided into two clusters based on the nasal expression levels of the top bronchial BE-associated genes. Participants in the cluster with the pattern of relative gene expression like the “bronchiectatic” cluster defined by bronchial gene expression had a higher radiologic BE burden, but we did not observe significant differences in clinical symptoms such as cough, phlegm, or dyspnea between the two clusters. This is in contrast to differences in symptom frequency between the participant clusters identified in Xu et al. based on bronchial BE-associated gene expression. In addition, an intermediate BE participant cluster had been identified in the prior bronchial BE study. Importantly, this intermediate cluster differed from the normal participant cluster by its higher proportion of current tobacco users (p < 0.01) but otherwise was noted to have similar clinical characteristics to that of the normal participant cluster, potentially highlighting the elevated risk of BE among those who smoke. In our analysis of the nasal validation dataset, we did not identify an intermediate BE cluster. One possible explanation for these findings is the lower burden of radiologic BE in the nasal dataset compared to the bronchial dataset. Alternatively, BE-related gene expression differences in nasal epithelium may not reflect these subgroup differences.

The enrichment of cell-type specific signatures derived from single-cell sequencing experiments among the genes whose expression is altered by radiologic BE in nasal epithelium suggests that similar changes in the proportion of deuterosomal, multiciliated, and basal cells occur in both the bronchial and nasal epithelium. These findings suggest that BE-associated changes in these cell populations extend into the upper airways. We previously hypothesized that an increase in multiciliated and deuterosomal cell-related gene expression and a decrease in basal cell-related gene expression in the bronchial epithelium might reflect a response to ciliated cell damage; if this hypothesis is correct, our present finding suggests that such damage extends into the upper airway.

One important limitation of the current study is that while it validates BE-related gene expression in independent samples, these validation samples are from the same cohort in which the bronchial BE-related gene expression signature was originally derived. Additional validation studies in independent cohorts are therefore required to demonstrate that these gene expression differences can be detected more broadly. It will be especially interesting to examine BE-related gene expression in individuals with a clinical diagnosis of BE, as well as individuals with less extensive history of tobacco smoke exposure.

Our study makes the novel observation that BE-related gene expression changes can be detected in nasal epithelium and validates our previously reported bronchial gene-expression signature of widespread radiologic BE in independent samples. Interestingly, while our data suggests that the impact of radiologic BE on nasal gene expression is less dramatic than it is for bronchial gene expression, we predict that radiologic BE alters the cellular composition of both compartments in a similar manner. As BE represents a cycle of epithelial dysfunction, chronic infection, recurrent inflammation, and structural damage, our bronchial-derived signature of BE may be applicable to other disease states with a similar pattern of chronic inflammation. Our validation results provide a potential avenue for non-invasive screening and disease monitoring in BE, as well as other disease patterns of chronic inflammation and airway dysfunction.

---

**Supplemental Table.** Clinical features of participants in the nasal validation set (n = 220)

|  | **No Widespread Radiologic BE (N = 204)** | **Widespread Radiologic BE (N= 16)** | **P-value** |
| --- | --- | --- | --- |
| Any Bronchiectasis (%) | 36 (18%) | 16 (100%) | <0.001 |
| Average number of lobes with BE (mean (SD)) | 0.27 (0.62) | 4.00 (1.15) | <0.001 |
| Male (%) | 142 (70%) | 12 (75%) | 0.87 |
| Race |  |  | 0.83 |
| Asian (%) | 5 (2.5%) | 0 (0%) |  |
| Black (%) | 37 (18.1%) | 4 (25%) |  |
| Others/Unknown (%) | 10 (4.9%) | 1 (6%) |  |
| White (%) | 152 (74.5%) | 11 (69%) |  |
| Smoking Status |  |  | 0.22 |
| Current tobacco user (%) | 68 (38%) | 3 (21%) |  |
| Former tobacco user (%) | 112 (62%) | 11 (79%) |  |
| FEV1 % predicted (mean (SD)) | 65 (18) | 81 (22) | 0.11 |
| FEV1/FVC (mean (SD)) | 0.59 (0.13) | 0.72 (0.15) | 0.026 |
| Cough Yes (%) | 93 (46%) | 8 (50%) | 0.97 |
| Phlegm Yes (%) | 92 (45%) | 7 (44%) | 0.53 |
| Shortness of Breath Yes (%) | 119 (58%) | 13 (81%) | 0.21 |
| DECAMP Cohort |  |  | 0.63 |
| DECAMP1 (%) | 77 (38%) | 7 (44%) |  |
| DECAMP2 (%) | 127 (62%) | 9 (56%) |  |

## References

1. Chalmers JD, Aliberti S, Blasi F. Management of bronchiectasis in adults. Eur Respir J. 2015;45(5):1446–1462.

2. Polverino E, Dimakou K, Hurst J, et al. The overlap between bronchiectasis and chronic airway diseases: state of the art and future directions. Eur Respir J. 2018;52(3).

3. Traversi L, Miravitlles M, Martinez-Garcia MA, et al. ROSE: radiology, obstruction, symptoms and exposure - a Delphi consensus definition of the association of COPD and bronchiectasis by the EMBARC Airways Working Group. ERJ Open Res. 2021;7(4).

4. Diaz AA, Nardelli P, Wang W, et al. Artificial Intelligence-based CT Assessment of Bronchiectasis: The COPDGene Study. Radiology. 2022:221109.

5. Diaz AA, Wang W, Orejas JL, et al. Suspected Bronchiectasis and Mortality in Adults With a History of Smoking Who Have Normal and Impaired Lung Function : A Cohort Study. Ann Intern Med. 2023;176(10):1340–1348.

6. King PT, Holdsworth SR, Freezer NJ, Villanueva E, Holmes PW. Characterisation of the onset and presenting clinical features of adult bronchiectasis. Respir Med. 2006;100(12):2183–2189.

7. Morrissey BM, Evans SJ. Severe bronchiectasis. Clin Rev Allergy Immunol. 2003;25(3):233–247.

8. Tiddens HAWM, Meerburg JJ, van der Eerden MM, Ciet P. The radiological diagnosis of bronchiectasis: what’s in a name? Eur Respir Rev. 2020;29(156).

9. Hill AT, Sullivan AL, Chalmers JD, et al. British Thoracic Society Guideline for bronchiectasis in adults. Thorax. 2019;74(Suppl 1):1–69.

10. Weycker D, Hansen GL, Seifer FD. Prevalence and incidence of noncystic fibrosis bronchiectasis among US adults in 2013. Chron Respir Dis. 2017;14(4):377–384.

11. ten Hacken NH, Wijkstra PJ, Kerstjens HA. Treatment of bronchiectasis in adults. BMJ. 2007;335(7629):1089–1093.

12. Beane J, Vick J, Schembri F, et al. Characterizing the impact of smoking and lung cancer on the airway transcriptome using RNA-Seq. Cancer Prev Res (Phila). 2011;4(6):803–817.

13. Billatos E, Vick JL, Lenburg ME, Spira AE. The Airway Transcriptome as a Biomarker for Early Lung Cancer Detection. Clin Cancer Res. 2018;24(13):2984–2992.

14. Gower AC, Steiling K, Brothers JF, Lenburg ME, Spira A. Transcriptomic studies of the airway field of injury associated with smoking-related lung disease. Proc Am Thorac Soc. 2011;8(2):173–179.

15. Steiling K, Lenburg ME, Spira A. Personalized management of chronic obstructive pulmonary disease via transcriptomic profiling of the airway and lung. Ann Am Thorac Soc. 2013;10 Suppl(Suppl):S190–196.

16. Campbell JD, McDonough JE, Zeskind JE, et al. A gene expression signature of emphysema-related lung destruction and its reversal by the tripeptide GHK. Genome Med. 2012;4(8):67.

17. Menon AA, Lee M, Ke X, et al. Bronchial epithelial gene expression and interstitial lung abnormalities. Respir Res. 2023;24(1):245.

18. Steiling K, Ryan J, Brody JS, Spira A. The field of tissue injury in the lung and airway. Cancer Prev Res (Phila). 2008;1(6):396–403.

19. Xu K, Diaz AA, Duan F, et al. Bronchial gene expression alterations associated with radiological bronchiectasis. Eur Respir J. 2023;61(1).

20. Billatos E, Duan F, Moses E, et al. Detection of early lung cancer among military personnel (DECAMP) consortium: study protocols. BMC Pulm Med. 2019;19(1):59.

21. Xu K, Diaz AA, Duan F, et al. Bronchial Gene Expression Alterations Associated with Radiographic Bronchiectasis. Eur Respir J. 2022.

22. Di Tommaso P, Chatzou M, Floden EW, Barja PP, Palumbo E, Notredame C. Nextflow enables reproducible computational workflows. Nat Biotechnol. 2017;35(4):316–319.

23. Dobin A, Davis CA, Schlesinger F, et al. STAR: ultrafast universal RNA-seq aligner. Bioinformatics. 2013;29(1):15–21.

24. Li B, Dewey CN. RSEM: accurate transcript quantification from RNA-Seq data with or without a reference genome. BMC Bioinformatics. 2011;12:323.

25. Xu K, Shi X, Husted C, et al. Smoking modulates different secretory subpopulations expressing SARS-CoV-2 entry genes in the nasal and bronchial airways. Sci Rep. 2022;12(1):18168.

26. Leek JT, Storey JD. Capturing heterogeneity in gene expression studies by surrogate variable analysis. PLoS Genet. 2007;3(9):1724–1735.

27. Wang L, Nie J, Sicotte H, et al. Measure transcript integrity using RNA-seq data. BMC Bioinformatics. 2016;17:58.

28. Ritchie ME, Phipson B, Wu D, et al. moderated powers differential expression analyses for RNA-sequencing and microarray studies. Nucleic Acids Res. 2015;43(7):e47.

29. Deprez M, Zaragosi LE, Truchi M, et al. A Single-Cell Atlas of the Human Healthy Airways. Am J Respir Crit Care Med. 2020;202(12):1636–1645.

30. Zhang X, Sebastiani P, Liu G, et al. Similarities and differences between smoking-related gene expression in nasal and bronchial epithelium. Physiol Genomics. 2010;41(1):1–8.

