## Supplemental Figure 1 for "Validation of bronchial airway gene expression associated with bronchiectasis in nasal epithelium"

**Supplemental Figure 1.** Schematic representation of data sources for nasal validation sample set.

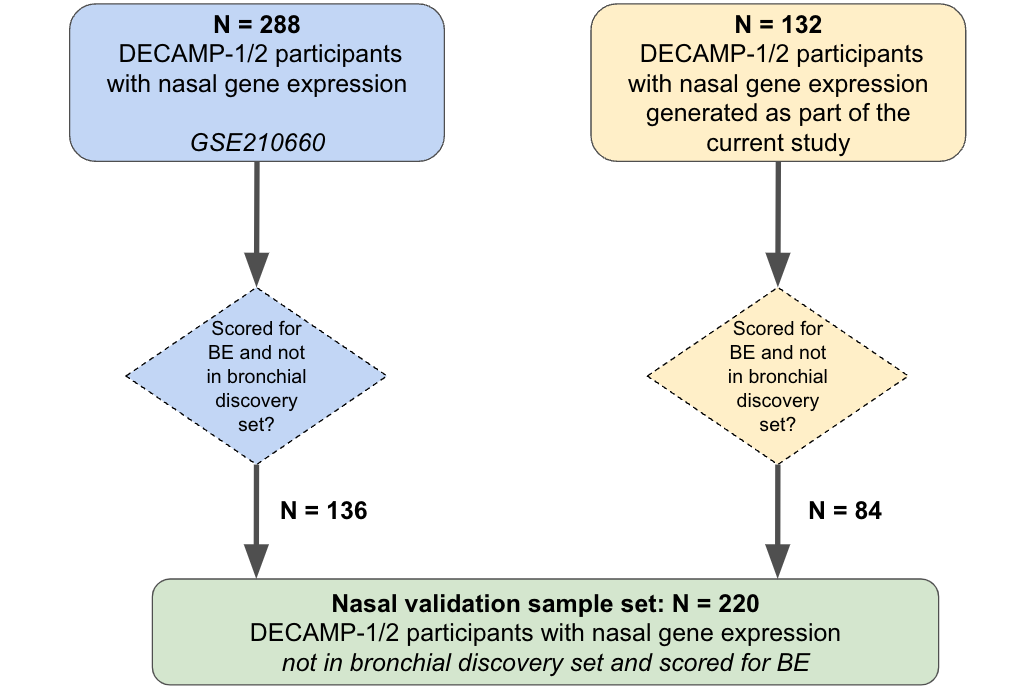
